# Effects of Task Demands on Neural Correlates of Acoustic and Semantic Processing in Challenging Listening Conditions

**DOI:** 10.1101/2021.01.22.427847

**Authors:** Dhatri S. Devaraju, Amy Kemp, David A. Eddins, Rahul Shrivastav, Bharath Chandrasekaran, Amanda Hampton Wray

**Author notes:** **Co-corresponding Authors:** Bharath Chandrasekaran, Department of Communication Sciences and Disorders, University of Pittsburgh, Pittsburgh, PA, Amanda Hampton Wray, Department of Communication Sciences and Disorders, University of Pittsburgh, Pittsburgh, PA.

## Abstract

**Purpose:** Listeners shift their listening strategies to prioritize lower-level acoustic information and higher-level semantic information in challenging listening conditions. However, the neural mechanisms underlying different strategies are unclear. The current study examined the extent to which encoding of lower-level acoustic cues is modulated by task demand and relationships with the higher-level semantic processing.

**Method:** Electroencephalography (EEG) was acquired while participants listened to sentences in noise that contained either higher or lower probability final words. Task difficulty was modulated by time available to process responses. Cortical tracking of speech - neural correlates of acoustic temporal envelope processing - were estimated using temporal response functions (TRFs).

**Results:** Task difficulty did not affect cortical tracking of temporal envelope of speech under challenging listening conditions. No correlations were observed between the cortical tracking of temporal envelope of speech and semantic processes, even after controlling for the effect of individualized signal-to-noise ratios.

**Conclusions:** Cortical tracking of temporal envelope of speech and semantic processing are differentially influenced by task difficulty. While increased task demands modulated higher-level semantic processing, cortical tracking of temporal envelope of speech may be influenced by task difficulty primarily when the demand is manipulated in terms of acoustic properties of the stimulus, consistent with an emerging perspective in speech perception.

## Introduction

In challenging conditions, listeners shift weighting of lower-level acoustic and higher-level semantic cues by adopting strategies to maximize perception. Increased reliance on semantic information was demonstrated under challenging conditions where the given time to process the response choices was altered (Kemp et al., 2019). Reducing the time given to process the response choices has been shown to increase the task demand (Benikos et al., 2013) and thereby alter semantic processing in challenging listening environments (Kemp et al., 2019). The study by Kemp and colleagues (2019) was designed to assess the influence of task demands on semantic encoding by leveraging well-studied electrophysiological measures of semantic processing. Participants listened to sentences with high- or low-probability final words in the presence of four-talker babble. Response options for the subsequent word identification task were presented visually for 400, 700, or 1000 ms. Reducing the time available to process the response options increased cognitive engagement for sentence processing, indexed by larger neural correlates of semantic processing for shorter response processing times.

Recent studies have demonstrated the ability to assess neural encoding of the temporal envelope of speech using a novel analytic approach (Di Liberto et al., 2015; Ding et al., 2014, 2016; Ding & Simon, 2014). Although not the original intent of Kemp et al. (2019), the data acquired provides a unique opportunity to assess the dynamic interactions between higher-level semantic and lower-level acoustic processing under task constraints. Reanalysis of EEG data from Kemp et al. (2019) was conducted in order to examine the impact of task demand (varying time to process the response choices in the presence of background noise) on the neural encoding of lower-level acoustic information as indexed by cross-correlation of the neural response to the temporal envelope of the acoustic speech stimulus. Prior studies have shown that the temporal envelope of speech, a complex pattern of amplitude modulations, is highly relevant to speech processing (Ding et al., 2014, 2016; Ding & Simon, 2014; Drullman, 1995; Drullman et al., 1994; Shannon et al., 1995, 1998; Smith et al., 2002). This analysis method has been used to demonstrate increased listening effort in difficult listening conditions, such as noisy speech (Decruy et al., 2020; Dimitrijevic et al., 2019; McHaney et al., 2020), non-native speech (Song & Iverson, 2018), and fast speech rate (Müller et al., 2019). The impact of allotted response time on listening effort under challenging listening conditions has not been assessed by indexing neural encoding of temporal envelope of speech, yet such a method can present the opportunity to reveal whether or not such demands impact the fidelity of the neural signal in response to speech. By comparison to the original analyses of Kemp et al. (2019), these new analyses offer the potential to evaluate the relationship between cortical representations of high-level semantic and low-level acoustic processing under the same conditions.

The TRACE model of speech perception (McClelland & Elman, 1986) and the Ganong effect (Ganong, 1980) emphasize the importance of lexical context in establishing acoustic approximations for deconstructing ambiguous sound or phonetic environment. Susceptibility to Ganong effect exhibits greater reliance on top-down semantic processing, which has shown to influence the bottom-up perception of speech in challenging listening conditions (Lam et al., 2017). Thus, it is plausible that the processing of lower-level acoustic cues, such as the temporal envelope, is influenced by lexical/semantic processing as a result of increased task demand in challenging listening conditions. Altering the time given to process the response choices has been shown to affect semantic processing (Kemp et al., 2019) mediated by greater top-down constraints resulting from task demand. However, it is unknown if top-down constraints imposed by increased task demand modulate the bottom-up encoding of acoustic landmarks in speech, such as acoustic temporal envelope of speech.

Neural/cortical tracking of the temporal envelope of speech has been used to study segmentation of speech (Ding et al., 2016; Ding & Simon, 2014), where the temporal envelope of speech is mapped on to the neural electroencephalography (EEG) to derive a correlation metric. The envelope to EEG mapping across various time lags yields the Temporal Response Function (TRF), which has peaks similar to the event-related potentials obtained for auditory stimuli (Fiedler et al., 2017, 2019). While a few studies suggest that cortical tracking of temporal envelope of speech changes with listening effort (Dimitrijevic et al., 2019; Song & Iverson, 2018) others do not (Decruy et al., 2020; Müller et al., 2019), indicating inconsistencies in the literature on the effects of task demand on cortical tracking of temporal envelope of speech. Thus, the aim of the current study is to determine differential impact of varying task demand (varying time given to process response choices) on bottom-up lower-level acoustic cues (i.e., cortical tracking of temporal envelope of speech) and its relationship with the top-down higher level semantic processing (i.e., N400 amplitudes) under challenging listening conditions (i.e., individualized signal-to-noise ratio). We hypothesized that with increasing task demands, listeners will weigh semantics more than acoustics, which will result in increased N400 amplitudes and reduced cortical tracking of temporal envelope of speech when decreased time is allotted to process response choices. Alternatively, listeners may not weigh acoustic processing differently based on task demand, resulting in modulation of semantic processing only.

## Method

### Participants

Electroencephalography (EEG) datasets from thirty adults (13 males & 17 females) in the age range of 19 to 39 years (mean age = 22.83 years, SD = 4.65 years) were included from Kemp et al. (2019). All participants were right-handed monolingual speakers of English with normal hearing (hearing thresholds below 20 dB HL from 500 Hz to 8 kHz; ANSI, 2010) and normal or corrected-to-normal vision (Snellen, 1862). The mean education level of the participants was partial college. Mean education level of their caregivers was also partial college, a proxy variable to participant’s childhood socio-economic status. A neuropsychological battery was administered to assess nonverbal intelligence using the Test of Nonverbal Intelligence – 4^th^ Edition (Brown et al., 2010), receptive and expressive language via the Listening and Speaking Vocabulary and Grammar subtests of the Test of Adolescent and Adult Language – 3^rd^ Edition (Hammill et al., 1994), and inhibitory executive function assessed by the Stroop: Color and Word Test (Golden et al., 2003). All participants performed within normative limits on each task (Kemp et al., 2019).

### Paradigm

Complete details of the experimental paradigm are available in Kemp et al. (2019). Briefly, EEG was recorded while the participants listened to a total of 300 English sentences (stimulus duration: mean = 3.44 seconds, min = 1.76 seconds, max = 6.12 seconds), half of which were high cloze probability sentences, where the last word had a high probability of occurrence (e.g., Joe watched TV sitting on the *<u>couch</u>*), and the other half were low cloze probability sentences, where the last word had less probability of occurrence (e.g., Joe watched TV sitting on the *<u>swings</u>*). Listening difficulty was increased with the addition of background babble. The babble level for each individual was determined prior to recording EEG as the level of four-talker babble that resulted in 70.7% correct sentence recognition performance. Babble was presented continuously at that level for each individual throughout EEG acquisition. Sentence stimuli were presented at 62 dB SPL, and the signal-to-noise ratios (SNR) varied from -0.5 to -8 dB across participants. This resulted in effortful conditions where all participants performed equally on speech recognition.

Participants listened to each sentence and responded by choosing the target word from a choice of four visually presented word choices (the three foils rhymed with the target word). The time window for which the choices were displayed on the screen was operationally termed the response time deadline (RTD). Listening effort was further manipulated by varying RTDs, adding to the task demand. RTD was either 400 ms (short), 700 ms (mid), or 1000 ms (long). The experiment was conducted in 12 blocks, with one RTD per block. Participants were informed about the RTD (short, middle, long) before starting each block in order to prepare them for the task demands in each block.

### EEG processing and analysis

EEG recorded from 32 electrodes (Biosemi Active-Two system, Netherlands) along with three ocular electrodes was pre-processed offline using EEGLAB v14.1.2 (Delorme & Makeig, 2004) in MATLAB R2019b (MathWorks Inc., Natick, Massachusetts, USA). EEG was re-referenced to the average of right and left mastoid electrodes. Re-referenced EEG was down sampled to 128 Hz. The resampled EEG was then band-pass filtered from 1-15 Hz using a non-casual FIR filter with hamming window having high pass and low pass filter order of 846 and 424 Hz, respectively. Spherical spline interpolation was used to substitute electrode activity when activity of a specific electrode across the entire time window exceeded the activity of the adjacent electrodes by more than three standard deviations. Artifact subspace reconstruction was performed using approximately 60 seconds of clean EEG segments (identified by manual inspection) as implemented in EEGLAB (Mullen et al., 2015) to remove artifacts, such as sudden bursts and drifts. The acoustic waveform for each sentence was aligned with the respective EEG epoch based on a window from -1 to 6 seconds time-locked to the onset of each sentence. Epochs with voltages greater than ±100µV were rejected. Independent component analysis was carried out on the epoched EEG to remove the ocular, EKG, and muscular artifacts.

### Estimation of Temporal Response Functions (TRFs)

Temporal response functions were obtained from multivariate linear ridge regression analysis using the mTRF toolbox v1.4 (Crosse et al., 2016). TRFs are regression coefficients as a function of neural latency. The regression coefficients quantify the relationship between the temporal speech envelope and the EEG. Higher absolute TRF magnitude indicates a stronger relationship between the speech acoustics and the EEG, thus indicating greater cortical tracking of the temporal envelope of speech (Crosse et al., 2016; Lalor & Foxe, 2010). TRFs were evaluated for encoding the first derivative of the temporal envelope of speech, which emphasizes the acoustic edges (phoneme/syllable onsets & offsets) in the speech envelope. The multiband speech temporal envelope was derived by filtering the sentence waveforms with a gammatone filter bank consisting of 16 filters spaced contiguously on an equivalent rectangular bandwidth (ERB) scale from 250 to 8000 Hz (Slaney, 1998). The absolute value of the temporal envelope at the output of each filter was obtained by applying the Hilbert transform. The absolute value at the output of each of the 16 bands was raised to a power of 0.6, which mimics the compression characteristics of the normal inner ear (Vanthornhout et al., 2018), which has the effect of reducing temporal envelope cues. It is known that the acoustic cues are distorted by the presence of simultaneous competing sounds (Cooke, 2006; Drullman, 1995). Given that onsets of the stimulus contribute more to the EEG signal than steady-state portions, studies investigating cortical tracking of the temporal envelope of speech in presence of noise have advocated the derivative method where the acoustic edges or onsets of the signal are enhanced using first-derivative of the temporal envelope (Fiedler et al., 2017, 2019).

TRFs were estimated by mapping the derivatives of speech temporal envelope onto the EEG using forward mapping. Forward mapping was done using leave-one-out cross validation (Crosse et al., 2016; Di Liberto et al., 2015; McHaney et al., 2020). Time lags from -100 to 450 ms were considered for TRF estimation. EEG signals obtained for all RTD conditions across all electrodes were used for selecting the ridge regularization parameter. The optimal ridge parameter between 2^0,1,2…20^ across conditions was estimated using cross validation. The similarity between EEG and the TRF predicted from the derivative of the stimulus temporal envelope was estimated using Pearson’s product moment correlation and considered as the cortical tracking measure (r). Cortical tracking measures were estimated using leave-one-out cross-validation, where the TRFs across all conditions were averaged to predict the EEG in the left-out trial. TRFs were trained by pooling across all RTD conditions since the total stimulus duration (summed across all sentences) within each RTD condition was too low to estimate TRFs. This process was reiterated to obtain the cortical tracking measure for every sentence presented. The cortical tracking measures across all electrodes for all sentences were separately averaged for each RTD condition. To examine if the cortical tracking measures were above-chance, cortical tracking for mismatched sentence and EEG pairs was performed using 1000 random permutations and the 95^th^ percentile of the distribution was determined as the chance level.

### N400 Mean Amplitudes

In Kemp et al. (2019), visual inspection of the data revealed that the N400 was most prominent over centroparietal electrode sites, consistent with previous studies (e.g., Kutas & Federmeier, 2011). Mean amplitudes were averaged across these electrodes (CP5, CP6, C3, C4, P7, P8, P3, P4, PO3, PO4, O1 & O2) within a typical N400 time window of 350 - 750 ms (see Kemp et al., 2019 for details). N400 mean amplitudes for low and high probability sentences for each RTD were considered separately.

### Statistical Analysis

The cortical tracking (r) of the temporal envelope and the N400 effect (difference in mean N400 amplitude between low cloze and high cloze probability sentences) across different RTDs were compared using repeated-measure ANOVAs. N400 mean amplitudes for two cloze (low and high) sentences across RTDs were compared using a separate repeated-measure ANOVA to establish correspondence between the current post-processing procedures and those used by Kemp et al. (2019). The TRFs obtained across three RTD conditions were compared using point-to-point-wise t-tests. The change in N400 mean amplitudes across the three RTD conditions (slope), separately for low and high cloze probability, and the change in cortical tracking (r) across the three RTD conditions (slope) were estimated by fitting a linear model. The relationships between the N400 mean amplitude and cortical tracking slopes were examined using Spearman’s Rank correlation coefficient. Spearman’s Rank correlation was used considering the lower number of data points and the variability among the data points. Partial correlations using Spearman’s Rank correlation coefficient were carried out to control for the effect of SNR. Alpha was set at p < 0.05.

## Results

### Temporal Response Functions

The temporal response functions (waveforms) derived across three different RTDs (400 ms, 700 ms, and 1000 ms) along with the scalp topographies (insets) are shown in Figure 1A. Temporal response functions show peaks at post stimulus lags, flat baselines and peak scalp topography consistent with the expected neural activity. The fronto-central scalp distribution of the cortical tracking of speech envelope (Figure 1B) is consistent with cortical auditory event-related potentials. Point-to-point-wise t-tests did not show significant differences among TRFs obtained for any RTD condition (p > 0.05). There was a significant and strongly positive correlation between cortical tracking of speech envelope among all the three RTD conditions [400 & 700 (ρ = 0.73, p = < 0.001), 400 & 1000 (ρ = 0.79, p = < 0.001), 700 & 1000 (ρ = 0.76, p = < 0.001); Figure 1D]. The relative stable cortical tracking of the speech temporal envelope derived across different RTDs is consistent with the lack of significant effect of RTD on cortical tracking [F(2,58) = 0.704, p = 0.499]. Figure 1B illustrates means and individual subject data. Seven out of thirty participants had cortical tracking values less than chance values. To ensure validity of these results, analyses were repeated without these participants. Figure 1C illustrates the individual cortical tracking values for participants whose TRFs were below chance levels. Excluding participants with below-chance cortical tracking values did not change results [RTD: F(2, 44) = 0.982, p = 0.383], further validating the findings. Therefore, these participants were included in correlational analyses.

**Figure 1.**
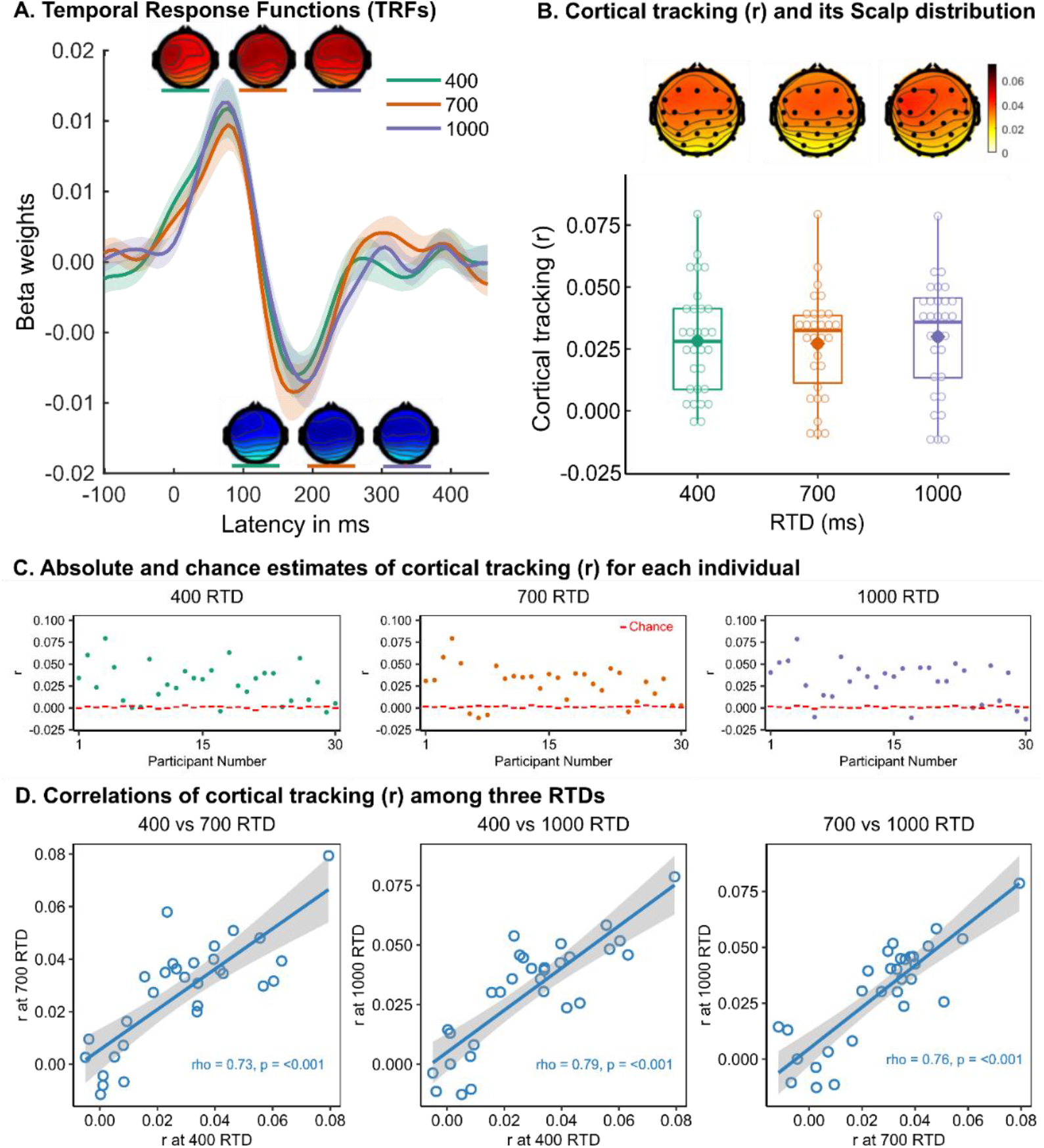
A. Mean temporal response functions (TRFs) in 400 ms, 700 ms and 1000 ms response time deadline (RTD) conditions averaged across all the electrodes. The shaded regions represent standard error of mean. The scalp topographies of the positive and negative peaks of TRFs for the three RTDs are shown above and below positive and negative peaks respectively. B. Boxplots of cortical tracking (r) values across 400 ms, 700 ms & 1000 ms RTD conditions averaged across all the channels. The middle line of the box represents median, top and bottom lines represents 25^th^ and 75^th^ percentile of the data, respectively. The dot inside the box represent the mean. The open circle represents individual participant data. Above the boxplot are scalp distributions of cortical tracking (r) values across three RTDs. C. Dot plot of cortical tracking (r) for each individual across 400 ms, 700 ms & 1000 ms RTD conditions. The red dashes for each subject indicate the 95^th^ percentile of the chance distribution. D. Scatterplots of the correlation among cortical tracking (r) at 400, 700 and 1000 ms RTD. The correlation coefficient and the p-value are provided for each of the scatterplots.

### Relationship between cortical tracking of speech envelope and semantic processing

Comparing N400 mean amplitudes for high and low cloze probability sentences across RTDs showed a significant effect of RTD [F(2,58) = 5.133, p = 0.013] and cloze [F(1,29) = 11.04, p = 0.02] and no interaction between cloze and RTD condition [F(2,58) = 0.115, p = 0.892], replicating the results of Kemp et al. (2019). Correlations between the change in, or slope of, N400 amplitudes between RTDs and the cortical tracking index between RTDs revealed no significant correlations between N400 amplitude slope and cortical tracking slope for either low probability (ρ = -0.07, p = 0.73) or high probability sentences (ρ = 0.07, p = 0.72). Additional partial correlational analyses between N400 amplitude and cortical tracking slopes controlling for the effect of SNR yielded similar results; no significant correlations for low (ρ = -0.06, p = 0.78) or high (ρ = 0.33, p = 0.13) probability sentences.

## Discussion

The current study evaluated differential processing of acoustic and semantic cues and their interplay during the varying task demands, i.e., changing response time deadlines under challenging listening condition, in the presence of four-talker babble. The effects and complements of two neural signatures (cortical tracking of temporal envelope of speech and the N400 response) during challenging listening conditions with varying task difficulty were assessed. Results indicate that cortical tracking of temporal envelope of speech and semantic processing are differentially affected by task difficulty. Cortical tracking of temporal envelope of speech remains stable across varying task difficulty while semantic processing varies as a function of change in task difficulty.

TRFs showed peaks that were consistent with previous literature (Ding & Simon, 2012; Fiedler et al., 2017, 2019; Müller et al., 2019). TRFs were similar across all RTDs and had similar scalp distributions, indicating no effect of increased task demand (i.e., shorter RTDs) on cortical tracking of temporal envelope of speech. These results are in line with Müller et al. (2019), who reported no correlation between neural tracking and listening effort as measured by subjective ratings and pupil dilation. Their task demands were varied using linguistic complexity (subject-verb-object occurrence) and speech rate (fast and slow). While linguistic complexity did not have any effect, speech rate did have a significant influence on neural tracking of temporal envelope of speech. The results of the present study, along with those of Müller et al. (2019), indicate that differences in the cortical tracking of temporal envelope of speech are evident only when the listening demands vary in terms of acoustics.

Further support for this pattern is found by Decruy et al. (2020), where cortical tracking of temporal envelope of speech varied across different levels of speech understanding but was not confounded by the listening effort. Song and Iverson (2018) reported similar findings, with differences in cortical tracking of temporal envelope of speech for conditions where acoustic properties varied, including stimuli presented in multiple languages by multiple speakers. Dimitrijevic et al. (2019) used stimuli manipulated with noise presented to adults with cochlear implants, also reporting differences in cortical tracking of temporal envelope of speech between conditions. Together with the current findings, these studies reveal that the cortical tracking of temporal envelope of speech appears to be influenced by task demands and/or listening effort only when the demand is manipulated in terms of acoustic properties of the stimulus, consistent with an emerging perspective.

Given that previous studies have shown the effect of SNR on cortical tracking of temporal envelope of speech (Decruy et al., 2020; Dimitrijevic et al., 2019; Lesenfants et al., 2019; Vanthornhout et al., 2018), a limitation of our experiment was that SNR was individualized. However, there were no correlations between semantic and acoustic processing across RTDs, even after controlling for the effects of SNR. The SNR did not show any relationship with the cortical tracking of temporal envelope of speech at any of the RTDs, which indicates that the acoustic cues were perceived and represented similarly, irrespective of the acoustic degradations when matched for speech understandability, and not modulated by task demand.

While neural tracking of temporal envelope of speech is not altered by RTDs, semantic processing was greater for the shorter RTD compared to longer RTD (Kemp et al., 2019). This indicates distinct neural processes for semantics and cortical tracking of acoustic temporal envelope that can be differentially manipulated in a single task and measured via EEG. Furthermore, semantic processing changes as a function of task demands do not appear to be associated with the cortical temporal envelope tracking. According to the TRACE model (McClelland & Elman, 1986), acoustic and phonetic word boundaries are filled in based on the semantic contexts, especially when acoustic cues are ambiguous, thus giving greater weight to semantic processing in challenging listening situations. The current findings support this model, indicating that listeners weigh higher-level semantic cues differentially depending on the listening condition. When higher level semantic cues are available, lower-level acoustic cues are engaged equally across listening conditions. The current results are in line with studies examining the effect of high and low cognitive load conditions on phoneme restoration in single word contexts, which suggest reduced reliance on acoustic cues in extracting the lexical output during high cognitive load conditions (Mattys et al., 2014; Mattys & Wiget, 2011). Higher-level semantic processing is dominant when listening difficulty is increased (e.g., short RTD or low cloze conditions), which requires higher cognitive resources, such as increased attentional resources. Reliance on acoustic cues is dominant only when the demand is exclusively bound to properties of the acoustic signal (e.g., timing, spectral characteristics). Taken together, the current findings suggest independence between domain-specific systems of audition and language in challenging listening environments, and that with increases in listening difficulty, listeners increasingly rely on semantic processing, without changing their reliance on the acoustic temporal envelope processing.

## Conclusions

The current study examined relationships between the two neural signatures, cortical tracking of temporal envelope of speech and N400, while listening to semantically altered sentences in the presence of a demanding task (varying RTDs) under challenging listening conditions (individualized SNR). While task difficulty modulated semantic processing, it did not modulate cortical tracking of temporal envelope of speech, indicating distinct processes for semantics and acoustics in challenging listening environments. Acoustic and semantic networks are domain-specific and may act independently of one other, with task demands inducing changes specific to the respective processes of each network. Top-down higher-level semantic processing may override bottom-up lower-level acoustic processing when the demanding task is not domain specific.

## Acknowledgements

We would like to acknowledge the support from the Michigan State University College of Communication Arts and Sciences Research Fellowship awarded to Amy Kemp, and the Richard and Nancy Heiss Endowment awarded to Amanda Hampton Wray. We are thankful to Nike Gnanateja for his support with analysis routines. Also, we would like to thank the participant pool in the study from Michigan State University.

